# Beyond conflict: kinship theory of intragenomic conflict predicts individual variation in altruistic behavior

**DOI:** 10.1101/2023.06.01.543237

**Authors:** Sean T. Bresnahan, David Galbraith, Rong Ma, Kate Anton, Juliana Rangel, Christina M. Grozinger

**Affiliations:** Department of Entomology, Center for Pollinator Research; Huck Institutes of the Life Sciences, Pennsylvania State University; Intercollege Graduate Degree Program in Molecular, Cellular, and Integrative Biosciences; Huck Institutes of the Life Sciences, Pennsylvania State University; Department of Entomology, Texas A&M University

## Abstract

Studies of the genetic basis of behavioral variation have emphasized gene cooperation within networks, often overlooking gene conflicts. The Kinship Theory of Intragenomic Conflict (KTIC) proposes that conflicts can occur within genes when parent-specific alleles have different strategies for maximizing reproductive fitness. Here, we test a prediction of the KTIC – that selection should favor alleles which promote “altruistic” behaviors that support the reproductive fitness of kin. In honey bee (*Apis mellifera*) colonies, workers act altruistically when tending to the queen by performing a “retinue” behavior, distributing the queen’s mandibular pheromone (QMP) throughout the hive. Workers exposed to QMP do not activate their ovaries, ensuring they care for the queen’s brood instead of competing to lay unfertilized eggs. Thus, the KTIC predicts that response to QMP should be favored by the maternal genome. Using a reciprocal cross design, we tested for parent-of-origin effects on the workers’ 1) responsiveness to QMP, 2) ovary activation, and 3) brain transcriptome. We hypothesized that QMP-responsive workers have smaller and less active ovaries, influenced by the workers’ parent-of-origin. With an allele-specific transcriptomic analysis, we tested whether QMP-responsive workers show enriched maternal allele-biased gene expression compared to QMP-unresponsive workers. Finally, we explored how parent-of-origin gene expression patterns are associated with overall gene expression patterns and regulatory networks. We report evidence in support of the KTIC for the retinue behavior and associated conflicts within gene networks. Our study provides new insights into the genetic basis of behavior and the potential for behavioral variation influenced by intragenomic conflict.

## Introduction

The Kinship Theory of Intragenomic Conflict (KTIC) provides an intriguing explanation for the often-observed variation in social behavior among individuals in family or social groups (Haig 2000), which is essential for adaptation and selection (Foster and Sih 2013). In social groups, behavioral variation generates flexible task repertoires and division of labor, which facilitate efficient collective responses to changing environments (Gordon 2014; Gordon 2016). Research into the molecular basis of individual behavioral variation has historically considered the cooperation between genes, particularly within gene networks (Jeanson and Weidenmüller 2014; Sinha et al. 2020). However, less attention has been given to conflicts that can occur within genes: as the KTIC proposes, conflicts can arise because parent-specific alleles have different strategies for maximizing their reproductive fitness (Gardner and Úbeda 2017). Previous studies of intragenomic conflict have focused on genes in which parent-specific alleles favor acting “selfishly,” increasing the reproductive fitness of the individual and reducing fitness of kin, such as in mammalian embryonic development and fetal growth (Barlow and Bartolomei 2014; Moore et al. 2015), seed production in plants (Rodrigues and Zilberman 2015), and reproduction in worker honey bees (Galbraith et al. 2016; Galbraith et al. 2021). The KTIC also proposes that parent-specific alleles can favor acting “altruistically,” reducing the individual’s reproductive output and promoting the reproductive fitness of kin. However, empirical tests of this hypothesis are still lacking.

According to the KTIC, genes within an individual’s genome can have conflicting interests, particularly regarding the allocation of maternal resources (Haig 1992; Haig 2000; Bourke 2014). Genes that promote altruistic behavior towards kin may be favored by natural selection even if they reduce an individual’s direct fitness. This is because altruism-promoting genes increase the individual’s inclusive fitness by increasing the likelihood of copies of those genes reproducing in the population. In contrast, genes that promote selfish behaviors that support direct fitness through reproduction should be selected to counteract the effects of altruism-promoting genes. This intragenomic conflict is predicted to occur frequently in species in which genes inherited from the mother (“matrigenes”) and the father (“patrigenes”) are not evenly distributed among the offspring genotypes (Gardner and Úbeda 2017). If females are polyandrous, their offspring share matrigenes but have different patrigenes. If their offspring use maternal resources for survival, matrigenes should be selected to act altruistically, reducing the use of maternal resources, and increasing cooperation between offspring, to ensure that sufficient resources are available for future offspring that carry these matrigenes. In contrast, patrigenes are predicted to act selfishly by maximizing the use of maternal resources and increasing competition between offspring, since the male may not be the father of future offspring (Patten et al. 2014).

The KTIC predicts that selection should favor a balance between the individual effects of genes engaged in conflict (Haig 2008; Patten et al. 2016), and so it is unclear whether or how they can contribute to physiology and behavior. However, such genes are frequently co-expressed (Al Adhami et al. 2015; Bonthuis et al. 2022), suggesting they may influence gene networks. Additionally, allelic differences in transcription factor (TF) occupancy are associated with allelic differences in transcription (Do Kim et al. 2006; McDaniell et al. 2010; Reddy et al. 2012). Therefore, it is hypothesized that small changes in expression of matrigenes and patrigenes can result in downstream effects on gene networks.

Honey bees are an outstanding model system for studying the causes and consequences of intragenomic conflict (Gibson et al. 2015; Kocher et al. 2015; Galbraith et al. 2016; Smith et al. 2020; Galbraith et al. 2021; Bresnahan et al. 2023). Due to their haplodiploid genetics, polyandrous mating system, and large colony size, matrigenes and patrigenes are unevenly distributed across colony members, and multiple social interactions are predicted to be influenced by intragenomic conflict (Queller 2003). Honey bee colonies have a single reproductive female queen, while her daughters (workers) typically remain sterile and forego their own reproduction to rear the queen’s offspring (Page 2013). If the queen is lost, workers can act selfishly, activating their ovaries and competing among each other to lay unfertilized eggs, or altruistically, by remaining sterile and rearing a new queen (Galbraith et al. 2015). Previous studies testing predictions of the KTIC in honey bees demonstrated that, as predicted, patrigenes favor the selfish behavior of worker ovary activation (Galbraith et al. 2016; Galbraith et al. 2021) and aggression (Bresnahan et al. 2023). However, those studies found that genes showing differences in parent-specific transcript abundance do not show differences in overall transcript abundance (Galbraith et al. 2016; Galbraith et al. 2021; Bresnahan et al. 2023). Thus, the question of how parent-specific gene expression leads to individual variation and fitness benefits for patrigenes remains to be determined.

Honey bees are also an excellent system in which to evaluate whether and how intragenomic conflict can lead to altruistic behavior. Worker bees act altruistically when they forgo their own personal reproductive fitness in favor of queen and sibling brood care (Ratnieks and Helanterä 2009). A pheromone produced by the queen—in particular, a five-component blend termed queen mandibular pheromone (QMP)—triggers behavioral and physiological changes in workers (Traynor et al. 2014; Grozinger 2015). Bees exposed to QMP show reduced worker ovary activation (Mumoki et al. 2018) and extended brood care activities (Pankiw et al. 1998). Moreover, QMP is a contact pheromone that releases a “retinue response” behavior in workers, attracting them to surround and groom the queen as she moves across the brood comb (Keeling et al. 2003). While tending to the queen in the retinue response, worker bees pick up QMP and spread it to other workers throughout the colony (Seeley 1979). Thus, the retinue response is an altruistic behavior, whereby the workers increase their exposure to QMP and spread it to the other bees, ensuring that all workers remain sterile and rear the queen’s offspring. According to the KTIC, the retinue response behavior should be favored by matrigenes (Queller 2003). Indeed, several studies have shown that there can be significant variation in retinue response behavior (Kocher et al. 2010; Galbraith et al. 2015), but the role of intragenomic conflict has not been investigated.

In this study, we tested predictions of the KTIC on the altruistic, pheromone-mediated retinue response behavior in worker bees. Using a reciprocal cross design, we assessed parent-of-origin effects on, and relationships between, responsiveness to QMP, ovary activation, and transcriptional profiles in the brain of worker bees. We tested the hypothesis that the physiological effects of QMP on workers (i.e., smaller, and less active ovaries) are influenced by a worker’s parent-of-origin. We conducted an allele-specific transcriptome analysis of QMP-responsive and QMP-unresponsive worker bees to test the hypothesis that a worker’s response to QMP is associated with an enrichment for maternally-biased gene expression. Finally, we explored how genes that show intragenomic conflict are associated with gene regulatory networks underlying behavioral variation.

## Methods

### Biological samples

Reciprocally crossed colonies were generated between F0 stocks of *A. mellifera ligustica* x *scutellata* hybrid honey bees (*i.e.*, “Africanized” honey bees, or “AHBs”) and European honey bees (“EHBs;” derived from *A. mellifera* subspecies that evolved in Europe), and between different stocks of EHBs. In studies conducted in 2019, we sourced reproductive females (queens) and males (drones) from two AHB colonies (colonies 1 & 2, or C1 & C2) and two EHB colonies of *A. m. ligustica* stock (C3 & C4); these were managed at the Janice and John G. Thomas Honey Bee Facility of Texas A&M University (TAMU) in Bryan, TX. In studies conducted in 2021, EHB queens and drones were sourced from four colonies of *A. m. ligustica* “Cordovan” stock (C5-8) (Koehnen & Sons, Inc., Glenn, CA), two colonies of a mixed *A. m. ligustica* stock (C9-C10) (BeeWeaver Honey Farm, Navasota, TX), one colony of *A. m. carnica* stock (C11), and one colony of *A. m. caucasia* stock (C12) (Old Sol Apiaries, Rogue River, Oregon); these were managed at a Penn State University apiary in University Park, PA. The source colonies were separated into blocks, with two F0 colonies from different genetic backgrounds assigned to each block (Figure 1).

**Figure 1.**
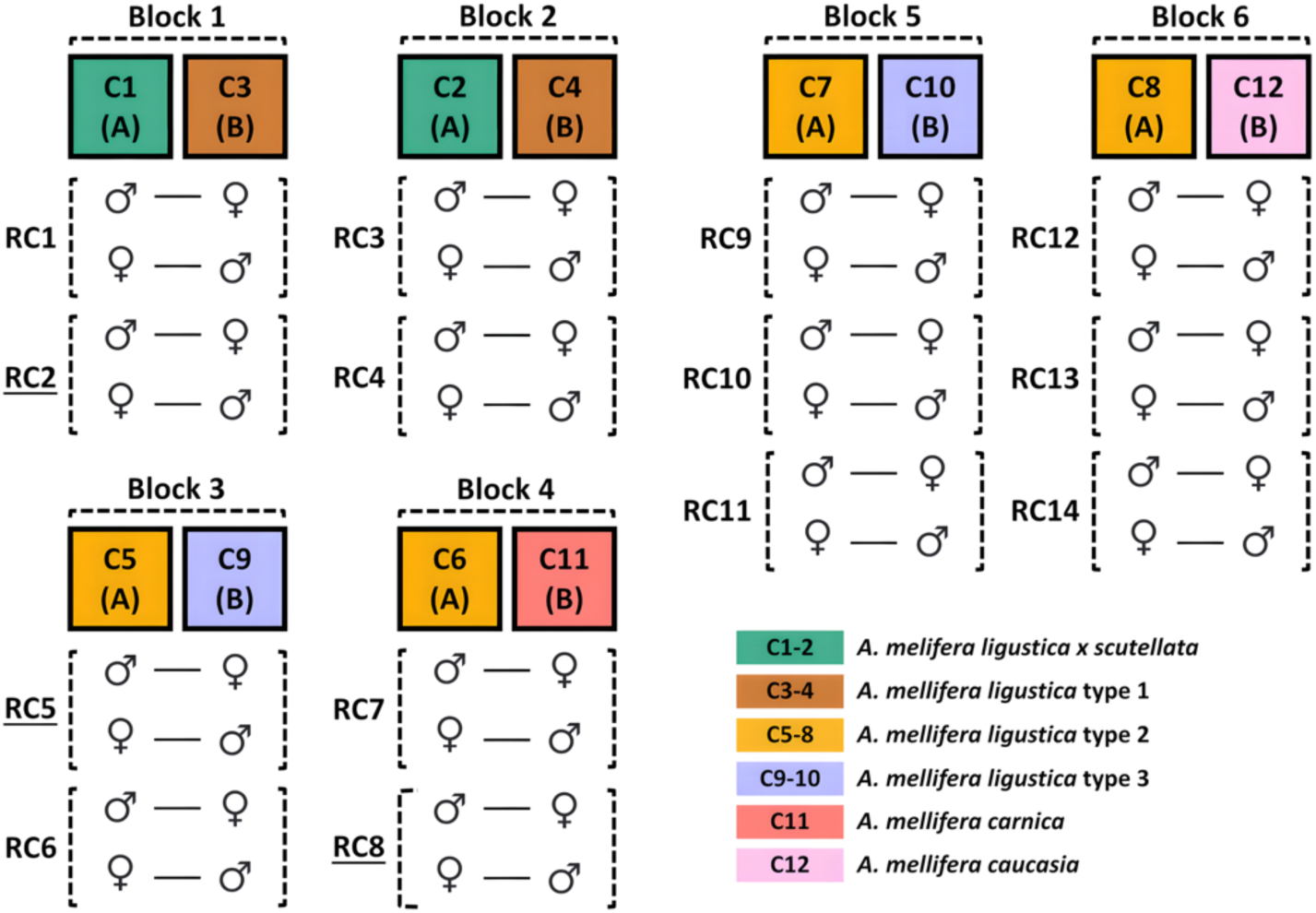
Reciprocal cross design. From each source colony (C1-C12), queens (♀) and drones (♂) were selected and crossed (AxB, BxA) to construct reciprocal crosses (RC1-RC14). Reciprocal crosses that were selected for sequencing are underlined. Colors indicate bee stock. Blocks 1-2 were generated for studies conducted in 2019, blocks 3-4 for studies in 2021, and blocks 5-6 for studies in 2022.

From each F0 colony, F1 queens were generated using a standard commercial practice known as grafting (Connor 2009). Adult virgin queens were labeled with a numbered tag affixed to the thorax and housed in cages within a nursery colony until F1 drones reared from these colonies reached sexual maturity. As drones tend to drift between colonies (Currie and Jay 1991), frames of emerging F1 drone brood were identified and isolated between queen excluders to prevent escape. Newly emerged F1 drones were marked on the thorax with non-toxic paint to identify their colony of origin and released back into the main colony entrance. In blocks 1-4, two F1 queens and two F1 drones were selected from each colony and crossed with two F1 queens and two F1 drones from the reciprocal colony by instrumental insemination (Cobey et al. 2013) to create two reciprocal crosses per block (reciprocal crosses 1-8, or RC1-8, Figure 1). In blocks 5-6, the same procedure was conducted with three F1 queens and three F1 drones from each colony to create three reciprocal crosses per block (RC9-14, Figure 1).

Inseminated queens were placed in their own colonies to initiate egg-laying, and the entrances of these colonies were restricted with queen excluder material to prevent escape or further mating. Approximately three weeks after the F1 queens began laying fertilized eggs, frames of sealed, emerging F2 brood were collected from each colony and stored in separate containers within a standing incubator kept at constant temperature of 34.5°C and 75% relative humidity. Once collection of the emerging F2 brood was completed, the queen from each colony was euthanized and stored at –80°C for DNA extraction and sequencing. The bodies of the drones used for inseminating each queen were likewise placed on dry ice and stored at –80°C immediately after semen collection for subsequent analysis.

In blocks 1-4, approximately five hundred age-matched newly emerged F2 worker bees from each F1 cross were collected on day 1 of the experiment and marked on the thorax with non-toxic paint to identify their maternal lineage of origin (Figure S1A). These bees were combined with an equal number of worker bees from the respective reciprocal crosses and placed into “nucleus” hive boxes containing honeycomb frames with honey and pollen but no queen or brood. In total, we created eight small nucleus colonies, each containing 500 F2 worker bees from each respective F1 cross in a reciprocal cross design. We also added 1,000 unmarked worker bees from an unrelated colony to each hive to increase the number of worker bees in the nucleus colony. Worker bees were kept in a “queenright” state (emulating exposure to the queen) by providing the colony with a plastic TempQueen strip (Mann Lake, Wilkes-Barre, PA) infused with QMP, as done previously (Pankiw et al. 1994).

In blocks 5-6, for each reciprocal cross, we populated six cages with five number-tagged age-matched newly emerged F2 worker bees from each F1 cross (Figure S1B). Workers were housed in upright petri dishes with pieces of beeswax foundation (Brushy Mountain Bee Farm, Moravian Falls, NC) fit against the back wall, following methods described in Shpigler and Robinson (2015). Each cage was provided a pollen ball at the bottom of the dish and a 50% sucrose solution via a modified 1.5 mL microcentrifuge tube for the bees to feed *ad libitum*. Worker bees were kept in a “queenright” state by providing a removable glass slide dosed with 0.1 queen equivalents of fresh QMP (Intko Supply LTD, Vancouver, BC) diluted in 1% water/isopropanol changed daily, following methods described in (Kocher et al. 2010).

### Parent-of-origin effects on the retinue response behavior

We assessed retinue response behavior in the F2 worker bees using two different methods. In blocks 1-4, we measured the colony-level frequency of retinue behavior as the proportion of individuals from each cross that was observed performing the retinue response (*i.e.*, touching with antennae and mouthparts) to the pheromone lure. On day 6 of the experiment, we affixed a fresh TempQueen strip to the center of an empty honeycomb frame (containing no food or brood) placed in the middle of each colony. Approximately 20 min after introducing the pheromone lure, we retrieved the frame. Over a ten-minute period, we collected paint-marked worker bees observed antennating the pheromone lure as being in a “responsive” behavioral state, and worker bees observed on the edges of the frame as being in an “unresponsive” behavioral state (Table S1). To test for an effect of maternal lineage (source colony) on the proportion of worker bees from each reciprocal cross colony that were observed performing the retinue behavior, we conducted logistic regression in R, treating the maternal lineage as a fixed effect nested within the block. We conducted this assay once for blocks 1-2 and three times on the same day spaced 3 hours apart for blocks 3-4 (Table S2).

In blocks 5-6, we measured the individual-level frequency of retinue response behavior as in Kocher et al. (2010). On days 4 to 6, we provided fresh QMP to each mini colony. Approximately 20 min after introducing the pheromone, we recorded responses to the stimulus (*i.e.* antennating and touching the slide where the QMP was added with their mouthparts) every 5 min for a total of 25 min, scoring a response as “1” and no response as “0” (Table S3). We used a linear mixed effects model, treating the cage as a random effect, to test for an effect of maternal lineage nested within the block on the retinue response scores of individual worker bees. Additionally, we performed one-way ANOVA tests to determine whether the frequency of retinue response across days varied between cages, between individual bees within cages, and/or between individual bees overall (Table S4).

### Parent-of-origin effects on ovary size and activation stage

The ovaries of ten individual worker bees per behavioral state, per cross, per reciprocal cross, and from the same blocks used for sequencing (10 bees x 2 behavioral states x 2 crosses x 2 reciprocally crossed colonies x 3 blocks = 240 individuals in total), were dissected under 1x phosphate buffered saline. Dissected ovaries were placed on a microscope slide and observed at 5x magnification for measurements of ovary size (the number of ovarioles present) and activation stage (four discrete stages or scores based on a modified Hess scale (Hess 1942) of 0 to 3 as done previously (Duncan et al. 2016) (Table S5). For each block, separately, one-way ANOVA tests were used to determine the significance of variation in ovary size and activation stage, respectively, by behavioral state and maternal lineage nested within behavioral state (Table S6). Additionally, one-way ANOVA tests were used to determine the significance of overall variation in ovary size and activation stage, respectively, by behavioral state and maternal lineage nested within block (Table S7).

### Sample preparation for whole genome re-sequencing and RNA-seq

The F1 queen and drone parents from each of three reciprocal crosses (RC2, RC5, RC8, underlined in Figure 1) constituting three distinct genetic blocks (blocks 1, 3, and 4) were selected for whole genome re-sequencing (WGS). The thorax of each bee was removed, and the flight muscle tissue was dissected on a platform surrounded by dry ice to keep the tissue frozen until the DNA isolation procedure. Genomic DNA was isolated using a DNeasy Blood & Tissue Kit (Qiagen, Germantown, MD). Samples were sent to the Penn State Genomics Core Facility for quantitation by a Qubit Fluorometer (Life Technologies, Carlsbad, CA) and quality control assessment by Genomic DNA ScreenTape analysis on a TapeStation 4150 (Agilent, Santa Clara, CA). Sequencing libraries were prepared using an Illumina DNA prep kit with unique 5’ and 3’ indexes, pooled, and sequenced on a NextSeq 2000 P3 flow cell to generate approximately 37 million 150-bp paired-end reads per sample (Table S8).

Three QMP-responsive and three QMP-unresponsive individuals of each maternal lineage (12 per 3 reciprocal crosses) were selected for sequencing. Heads were detached, and whole brains were dissected on dry ice and placed immediately in TRIzol (Invitrogen). Total RNA was isolated via TRIzol/chloroform extraction followed by precipitation with isopropanol. Samples were submitted to the Penn State Genomics Core for quantitation by Qubit and quality control assessment by RNA ScreenTape analysis. In total, 36 sequencing libraries were prepared using Illumina TruSeq Stranded mRNA Prep kits with unique 5’ and 3’ indexes, pooled, and sequenced on a NextSeq 2000 P3 flow cell to generate approximately 34 million 150-bp single-end reads per sample (Table S8). WGS and RNA-seq reads for this study are deposited in the NCBI Sequence Read Archive under SRA project accession PRJNA732718.

### Generation of parent genomes and identification of SNPs within transcripts

WGS reads were adapter trimmed with fastp (Chen et al. 2018), then aligned to the *A. mellifera* Amel_HAv3.1 genome assembly (Wallberg et al. 2019) with BWA-MEM (Li 2013) using the default settings. Alignment files were coordinate sorted using samtools (Danecek et al. 2021). SNPs were detected using freebayes (Garrison and Marth 2012) to account for differences in ploidy between queens and drones. Heterozygous SNPs were filtered using bcftools (Danecek et al. 2021) and homozygous SNPs were integrated into the Amel_HAv3.1 genome assembly with gatk (McKenna et al. 2010). Variant call files containing homozygous SNPs were intersected using bedtools (Quinlan and Hall 2010) to identify SNPs that were unique to each parent but shared between the crosses of a reciprocal cross pair. Parent-specific homozygous SNPs were then intersected with the RefSeq *A. mellifera* genome feature file to annotate SNPs within the longest transcript for each gene. SNPs intersecting more than two transcripts or two transcripts annotated on the same strand were filtered to reduce ambiguity in read alignments to multiple features. Additionally, transcripts for miRNA and tRNA genes were filtered, as these loci are inappropriate for allele-specific expression analysis using short sequencing reads (Wang and Clark 2014). This left 11,865 of 12,318 (96.3%) of the *A. mellifera* genes for analysis.

### Detection of allele-specific gene expression

RNA-seq reads were adapter trimmed with fastp (Chen et al. 2018), then aligned to each respective parent genome using tophat2 (Kim et al. 2013) with the settings “--library-type fr-firststrand --b2-very-sensitive.” Alignments with mismatches and secondary alignments were filtered using bamtools (Barnett et al. 2011) with the settings “-tag XM:0 -isPrimaryAlignment true.” For each sample, read coverage was calculated at each SNP using bedtools intersect in strand-aware mode. Coverage files were then combined in R to create a matrix of F2 read counts at F1 SNPs within *A. mellifera* transcripts. Whenever two or more SNPs received the exact same counts across all samples within a transcript (because the distance between each SNP was shorter than the length of the reads which aligned to them), one SNP was chosen at random. Additionally, SNPs that received less than 10 counts were filtered out. Finally, transcripts with less than 2 SNPs were filtered out prior to downstream analyses. Allele-specific counts of F2 RNA-seq reads at F1 SNPs within transcripts are available in Tables S9-S11.

We utilized the Storer-Kim (SK) binomial exact test of two proportions (Storer and Kim 1990; Wang and Clark 2014) and a general linear mixed effects model with interaction terms (GLIMMIX) to identify transcripts that exhibited parent-of-origin expression biases following methods described previously (Kocher et al. 2015; Galbraith et al. 2016; Galbraith et al. 2021; Bresnahan et al. 2023), with a few modifications. To adjust for variation in sequencing depth, we 1) fit the GLIMMIX models to the raw read counts and specified library size factors as an offset term, and 2) used normalized read counts for the SK tests. Library size factors were estimated using the median of ratios normalization (MRN) method implemented in the DESeq2 package (Love et al. 2014) on the within-sample SNP-wise sums of allelic read counts (Castel et al. 2015; Fan et al. 2021), treating maternal lineage and behavioral state as cofactors. Normalized read counts were generated by dividing the raw read counts by these size factors.

SK tests were conducted for each transcript for each SNP to test for significant differences in the proportion of maternal and paternal read counts (Tables S12-S14). Test results were then corrected to control for the false discovery rate (FDR) (Benjamini and Hochberg 1995) and aggregated by transcript. For each transcript, all positions were required to exhibit the same direction of parent- or lineage-specific bias at previously established thresholds (Wang and Clark 2014) for *p1* (the proportion of reads mapping to the “A” allele in offspring from the “B” queen and “A” drone), and *p2* (the proportion of reads mapping to the “A” allele in offspring from the “A” queen and “B” drone) for the transcript to be reported as exhibiting bias. Additionally, a GLIMMIX was fit for each transcript to test for an effect of parent, maternal lineage, and their interaction, offset by the log of the library size factors, and treating SNP and individual as random effects (Tables S15-S17). Test results were FDR corrected, and only transcripts with a significant effect of parent or lineage, but not their interaction, were considered to exhibit bias. Only genes considered to exhibit bias in both tests were reported as exhibiting bias (Table S19). Chi-squared tests were performed to determine whether there was an enrichment for any category of allelic expression bias (paternal, maternal, lineage A, lineage B) in QMP-responsive relative to QMP-unresponsive bees.

### Differential gene expression analysis

We used Kallisto (Bray et al. 2016) to estimate transcript abundances. A transcriptome index was generated with gffread (Pertea and Pertea 2020) using the Amel_HAv3.1 RefSeq genome feature file. Transcript abundance files were combined to generate a gene expression matrix (Table S18) using the tximport package (Soneson et al. 2022) in R for analysis of differential expression with DESeq2 (Love et al. 2014). After identifying and removing an outlier sample using a PCA projection pursuit method (Chen et al. 2020) (Figure S6), we employed a two-factor design to test for differential expression between behavioral states, treating maternal lineage as a cofactor (Table S20).

### Support Vector Machine (SVM) classification

SVM classification was conducted using the transcriptomes of QMP-unresponsive bees (coded with a value of 0) and QMP-responsive bees (coded with a value of 1). Transcriptomes for SVM classification were normalized to adjust for variation in sequencing depth between libraries by variance stabilizing transformation followed by quantile normalization. Genes with low counts and zero variance between samples were filtered from the dataset, leaving 11,514 of the 12,123 genes (94.98%) for SVM training. We subset the transcriptomic data on a random selection of samples into 80% training and 20% testing sets. Classifiers were fit using a radial kernel function, and the hyperparameters cost (*C*) and gamma (*γ*) were tuned via grid search in a parameter space of 2^-2^ < *C* < 1 and 10^-3^ < *γ* < 10^-1^. Feature selection was performed following methods described in Taylor et al. (2021). Starting with a classifier fit with all 11,514 genes, we iteratively performed the following: (1) the mean threefold cross-validation error of 20 rounds using random bins was calculated; (2) the matrix product of the classifier’s coefficients and its support vectors was used to calculate the feature weights of all remaining genes; (3) the gene with the smallest absolute weight was removed, until only 100 genes (an arbitrary minimum number of features chosen *a priori*) remained. The support vector classifier with the minimum cross-validation error or minimum number of features was then selected for downstream analyses (Figure S7; Table S21).

### Transcription regulatory network inference

Weighted Gene Co-expression Network Analysis (WGCNA) was performed using the WGCNA package (Langfelder and Horvath 2008) in R. Signed co-expression network modules were constructed using a soft-threshold power of 15 to balance scale independence and mean connectivity (Figure S8). Modules with eigengene correlation coefficients > 0.75 were merged (Figure S9; Table S22). We then calculated correlation coefficients of each module with behavioral state. Hypergeometric tests were conducted to identify modules enriched for genes with parent-of-origin expression biases and genes in our minimal support vector classifier of behavioral state.

The Analyzing Subsets of Transcriptional Regulators Influencing eXpression (ASTRIX) method for transcription regulatory network (TRN) inference (Chandrasekaran et al. 2011; Shpigler et al. 2017; Shpigler et al. 2019; Jones et al. 2020) was performed in MATLAB. Lists of honey bee orthologs of *Drosophila melanogaster* transcription factor (TF) genes were retrieved from Kapheim et al. (2015) and Jones et al. (2020). TFs predicted to interact with a target gene by ASTRIX were required to fulfill three criteria: (1) share a significant degree of mutual information with the target gene (𝑝 < 1 × 10^−6^), (2) explain at least 10% of the variance of the target gene, quantified by least angle regression, and (3) be predicted with a correlation of at least 0.8 using expression levels of the TF and target gene (Table S23). We then subset the TRN to include only those genes predicted to interact with parent-of-origin biased TFs. With this subset of the TRN, we performed hypergeometric tests to identify gene co-expression network modules enriched for the targets of parent-of-origin biased TFs.

### Gene Ontology (GO) enrichment analyses

GO enrichment analyses were conducted with DAVID (Sherman et al. 2022) using *D. melanogaster* orthologs of the genes showing parent-of-origin expression biases (Table S24), the genes in our minimal support vector classifier of behavioral state (Table S25), and the gene co-expression network modules constructed with WGCNA (Table S26). *Drosophila melanogaster* orthologs of *A. mellifera* genes were retrieved from Bresnahan et al. (2022). For this study, we focused on GO terms (specifically, biological processes and molecular functions) and KEGG pathways found to be significantly overrepresented among candidate genes at a false discovery rate of < 0.05.

## Results

### Parent-of-origin effects on the retinue behavior

Our results suggest a parent-of-origin effect on – and considerable individual variation in – the retinue response behavior in worker bees. We observed a significant association between maternal lineage and the proportion of worker bees performing the retinue response behavior in blocks 3 and 4, but not in blocks 1 and 2 (Figure 2A; Table S2). Additionally, in blocks 5 and 6, we observed a significant association between maternal lineage and the frequency of retinue response across days of individual bees (Figure 2B), between individual bees (Figure 2C), between cages (Figure S2), and between individual bees within cages (Table S4).

**Figure 2.**
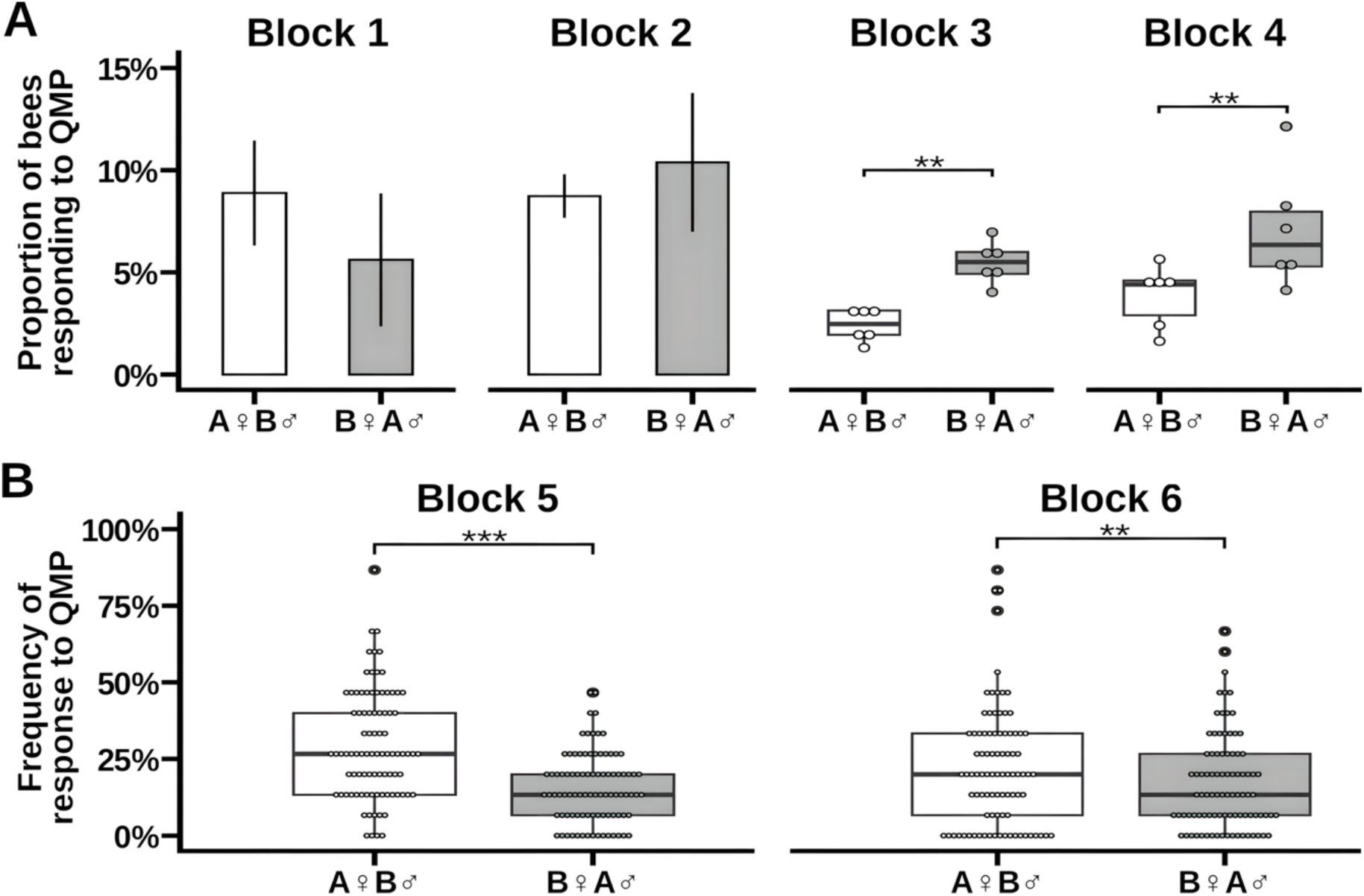
Parent-of-origin effects on the retinue response behavior. **(A)** Blocks 1-4, the proportion of bees responding to QMP of each maternal lineage (source colony) from two reciprocal crosses per block. Significance of one-way ANOVAs are indicated (** = p < 0.01). **(B)** Blocks 5 and 6, the mean frequency of response to QMP of individual bees, indicated by points, from six cage colonies constructed from each of three reciprocal crosses per block. Mixed-effect model significance of the effect of maternal lineage nested within blocks are indicated (** = p < 0.01, *** = p < 0.001).

### Parent-of-origin effects on ovary activation

To test for an association between maternal lineage and reproductive physiology, we evaluated ovary size and activation in blocks 1, 3, and 4, which included the reciprocal crosses that we subsequently used for sequencing. Bees in blocks 5 and 6 were not used for the remainder of our study as they were collected as older nurses for a separate study. We found that workers from colonies showing more frequent responses to QMP had smaller ovaries in all tested blocks, but only those in blocks 3 and 4 showed a significant effect of maternal lineage. Specifically, individuals from the A♀B♂ cross in blocks 3 and 4 had larger ovaries (Figure 3A, Table S6) – notably, these individuals also responded less frequently to QMP (Figure 2A). In a full model including all three blocks, ovary size varied significantly by behavioral state and maternal lineage (Table S7). These data suggest there is a parent-of-origin effect on ovary size and there are larger ovaries in QMP-unresponsive compared to QMP-responsive bees.

**Figure 3.**
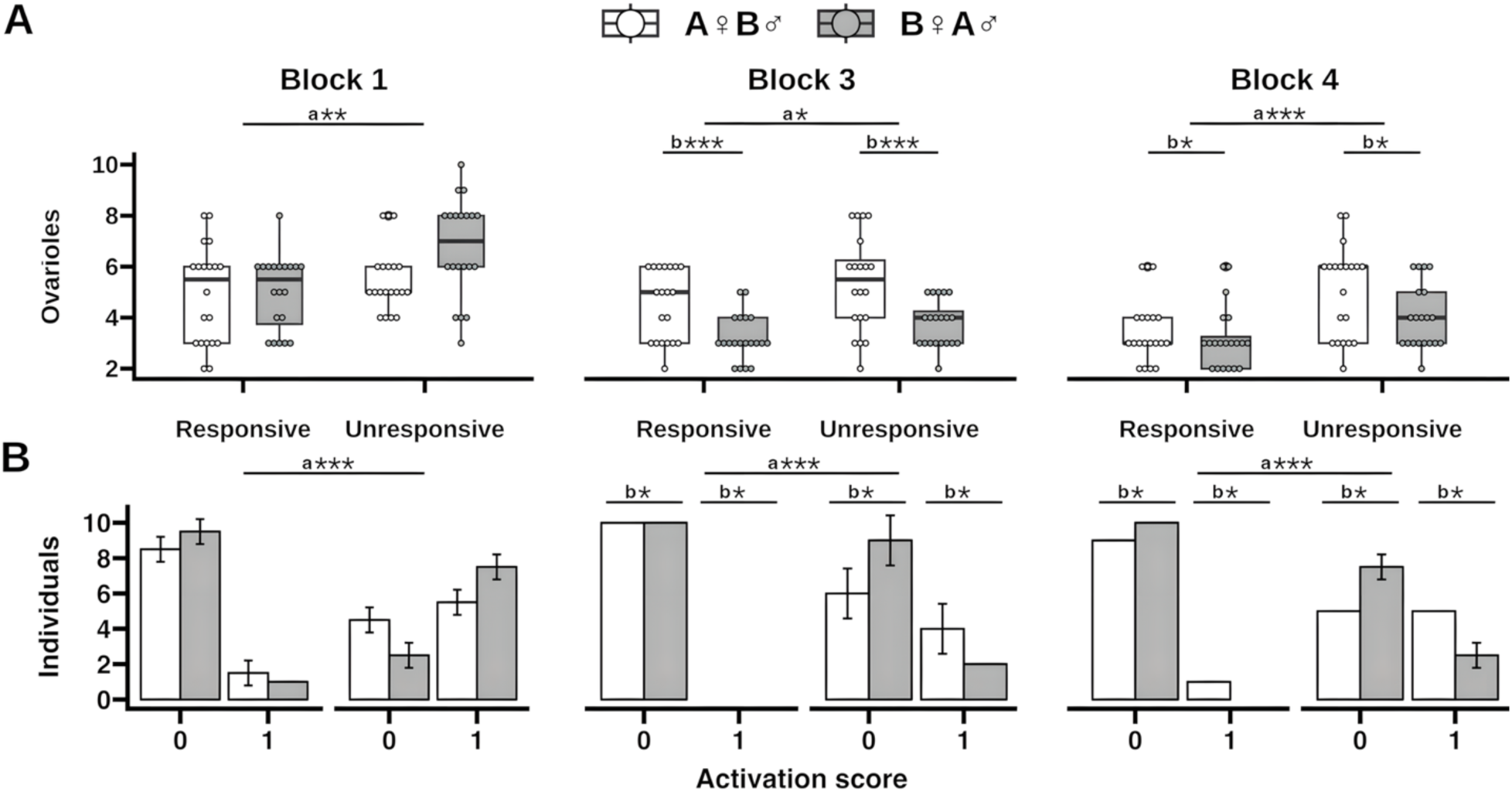
Parent-of-origin effect on ovary size and activation. The abdomens of 10 individuals of each behavioral state of each maternal lineage (source colony) from two reciprocal crosses per block were dissected to quantify **(A)** ovary size, and **(B)** ovary activation (0 to 3). Note: no ovaries at stages 2 and 3 were present in our samples. One-way ANOVA test significance of behavioral state (a) and maternal lineage (b) are indicated (* = *p* < 0.05, ** = *p* < 0.01, *** = *p* < 0.001).

Ovary activation varied significantly by behavioral state in all tested blocks, and by maternal lineage in blocks 3 and 4 but not in block 1 (Figure 3B, Table S6). Specifically, in all three blocks, ovaries were more active in QMP-unresponsive compared to QMP-responsive bees, and ovaries were more active in individuals from the A♀B♂ cross in blocks 3 and 4. In a full model including all three blocks, ovary activation varied significantly by behavioral state and maternal lineage (Table S7). These data suggest there is a parent-of-origin effect on ovary activation, and bees that did not respond to QMP had more active ovaries. Taken together, we found that bees from the maternal lineage that responded less frequently to QMP had larger and more active ovaries.

### Parent-of-origin gene expression

At approximately 40x genome coverage, we detected an average of 2.77 million homozygous SNPs per F1 diploid queen and 2.87 million SNPs per F1 haploid drone. On average, 550,695 of these SNPs were within transcripts and varied in their sequence identity between the parents of each cross, allowing for identification of parent-of-origin reads in the F2 worker bees, 491,338 (89%) of which were within transcripts that had at least two SNPs. After filtering SNP positions with low RNA-seq read coverage in the F2 bees, our datasets contained read coverage at an average of 77,131 positions distributed among an average of 4,366 of 11,865 transcripts (36.8%) in 12 QMP-unresponsive bees per block, and an average of 78,063 positions distributed among an average of 4,395 of 11,865 transcripts (37%) in 12 QMP-responsive bees per block (Tables S9-S11). Considering the overlap between behavioral states, on average, bees in each block expressed 4,143 of 11,865 transcripts (34.92%).

Accounting for all unique transcripts across the three blocks, we identified 624 maternally-biased transcripts in whole brains of unresponsive bees and 887 in responsive bees, in addition to 143 paternally-biased transcripts in unresponsive bees and 181 in responsive bees (Table S19). Of these transcripts, 64 showed consistent allele-biased expression in at least 2 blocks, all of which were biased toward the maternal allele. We identified 542 transcripts that showed allelic bias in block 1, 616 in block 3, and 609 in block 4. Some transcripts showed allelic bias in both unresponsive and responsive individuals (82 in block 1, 108 in block 3, 131 in block 4), whereas others showed allelic bias in only one behavioral state (unresponsive only: 184 in block 1, 191 in block 3, 178 in block 4; responsive only: 276 in block 1, 317 in block 3, 300 in block 4). In support of our hypothesis, we found that maternally-biased transcripts were enriched in QMP-responsive worker bees relative to QMP-unresponsive bees in all three blocks (see Figure 4 for results from block 3 and Figures S3-S4 for results from blocks 1 and 4). Parent-biased transcripts in responsive bees were enriched for genes encoding proteins with chromatin binding function, including several core components of polycomb and trithorax group complexes. Overall, parent-biased genes were enriched for protein and sequence-specific DNA binding (Table S24).

**Figure 4.**
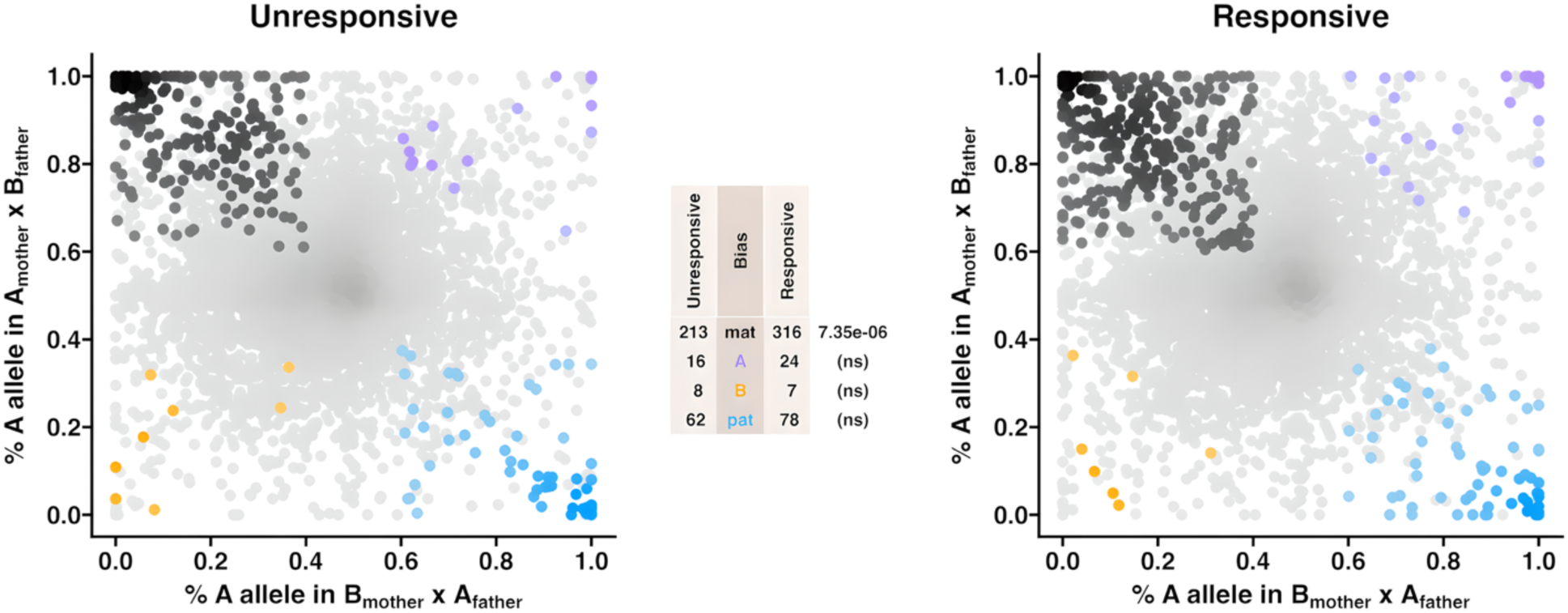
Worker bee response to QMP is associated with maternal allele-biased transcription. Allele-specific transcriptomes were assessed in worker bees from a reciprocal cross between different stocks of EHBs (C5xC9, block 3) that were unresponsive or responsive to QMP. The x-axis represents, for each transcript, the proportion of cross A (C9 lineage) reads in bees with a cross B (C5 lineage) mother and cross A father (*p1*). The y-axis represents, for each transcript, the proportion of cross A reads in bees with a cross A mother and cross B father (*p2*). Each color represents a transcript which is significantly biased at all tested SNP positions: black is maternal, purple is cross A, gold is cross B, blue is paternal, and gray is not significant. Significance was determined using the overlap between two statistical tests: a generalized linear interactive mixed model (GLIMMIX), and a Storer-Kim test along with previously established cutoff thresholds of *p1* < 0.4 and *p2* > 0.6 for maternal bias, *p1* > 0.6 and *p2* < 0.4 for paternal bias, *p1* < 0.4 and *p2* < 0.4 for lineage B bias, and *p1* > 0.6 and *p2* > 0.6 for lineage A bias. Similar results were obtained in blocks 1 and 4, see Figures S3-S4 for details.

### Gene expression differences

In both a full model accounting for all three blocks and a reduced model excluding the blocking factor, we did not detect any significantly differentially expressed genes in whole brains of QMP-unresponsive worker bees compared to QMP-responsive bees (Table S20). We therefore used support vector classification to identify more subtle gene expression differences which best classify individuals into behavioral states (Taylor et al. 2021). We trained a SVM using transcriptomes of 14 QMP-responsive bees and 15 QMP-unresponsive bees. A full model based on all 11,514 genes which passed our low count threshold achieved a root mean squared validation error of 0.5105 in threefold cross-validation. To identify model fit with a minimal set of genes that were maximally predictive of behavioral state, we conducted feature selection to progressively remove uninformative genes. The minimal model, which was trained on 100 genes (Table S21), had a root mean squared classification error of 0.2093, which was a significant improvement over that achieved by the full model (Figure S5). Genes in the minimal model were enriched for 15 GO terms including negative regulation of histone modifications, negative regulation of DNA recombination, nucleosome assembly, and chromosome organization (Table S25). Parent-of-origin biased genes were not significantly overrepresented in the model (8/100, *p* = 0.94).

### Inference of regulatory networks

We performed weighted gene co-expression network analysis (WGCNA) to identify modules of co-expressed genes (Langfelder and Horvath 2008). Additionally, we used the Analyzing Subsets of Transcriptional Regulators Influencing eXpression (ASTRIX) method (Chandrasekaran et al. 2011; Shpigler et al. 2017; Shpigler et al. 2019; Jones et al. 2020) to determine whether intragenomic conflicts within these co-expressed gene modules are influenced by transcription factors with parent-of-origin expression biases. For both analyses, we used the same 11,514 genes adjusted by variance stabilizing transformation and quantile normalization as in support vector classification.

Using this approach, we identified 22 co-expressed gene modules (Table S22) and inferred a transcription regulatory network consisting of 4,200 interactions between 196 transcription factors and 1,839 target genes (Table S23). Within this network, 29 of the 196 transcription factors (14.8%) showed parent-of-origin expression biases and were predicted to interact with 353 of the 1,839 target genes (19.2%). Specific co-expression modules were 1) correlated with behavioral state, 2) overrepresented for parent-biased genes, 3) overrepresented for genes in our minimal support vector classifier of behavioral state, and 4) overrepresented for the targets of parent-biased TFs (Table 1).

**Table 1.** Gene co-expression network modules are significantly correlated with behavioral state, and enriched with parent-biased genes, behavior-state classifying genes, and targets of parent-biased TFs.

| Module (size) | Correlation coefficient with behavioral state (*p*-value) | Parent-biased genes in module (enrichment *p*-value) | SVM genes in module (enrichment *p*-value) | Targets of parent-biased TFs in module (enrichment *p*-value) |
| --- | --- | --- | --- | --- |
| Module 1 (108) | -0.06 (0.70) | 24 (2.67e-03) | 1 (0.60) | 0 (1.00) |
| Module 2 (1868) | 0.14 (0.40) | 272 (8.78e-04) | 17 (0.40) | 148 (7.76e-43) |
| Module 3 (1518) | 0.27 (0.10) | 231 (1.85e-04) | 11 (0.74) | 67 (3.16e-05) |
| Module 4 (154) | 0.34 (0.04) | 34 (4.50e-04) | 1 (0.73) | 6 (0.20) |
| Module 5 (63) | 0.28 (0.10) | 11 (0.15) | 0 (1.00) | 0 (1.00) |
| Module 6 (84) | 0.19 (0.30) | 15 (0.09) | 0 (1.00) | 0 (1.00) |
| Module 7 (103) | 0.37 (0.03) | 30 (3.85e-06) | 1 (0.58) | 2 (0.74) |
| Module 8 (317) | 0.25 (0.10) | 57 (1.94e-03) | 2 (0.75) | 0 (1.00) |
| Module 9 (614) | 0.08 (0.70) | 67 (0.88) | 18 (2.96e-06) | 1 (1.00) |
| Module 10 (788) | 0.27 (0.10) | 94 (0.65) | 16 (8.18e-04) | 36 (4.37e-04) |
| Module 11 (74) | 0.11 (0.50) | 8 (0.71) | 1 (0.47) | 0 (1.00) |
| Module 12 (33) | -0.01 (0.90) | 6 (0.21) | 0 (1.00) | 0 (1.00) |
| Module 13 (162) | -0.03 (0.90) | 19 (0.63) | 0 (1.00) | 3 (0.78) |
| Module 14 (216) | 0.01 (1.00) | 31 (0.21) | 2 (0.54) | 4 (0.80) |
| Module 15 (74) | -0.19 (0.30) | 0 (1.00) | 0 (1.00) | 2 (0.57) |
| Module 16 (112) | -0.20 (0.30) | 13 (0.63) | 1 (0.61) | 0 (1.00) |
| Module 17 (739) | -0.19 (0.30) | 50 (1.00) | 4 (0.88) | 0 (1.00) |
| Module 18 (361) | 0.00 (1.00) | 21 (1.00) | 1 (0.95) | 0 (1.00) |
| Module 19 (106) | -0.13 (0.50) | 8 (0.96) | 0 (1.00) | 0 (1.00) |
| Module 20 (1963) | -0.33 (0.05) | 157 (1.00) | 14 (0.79) | 15 (1.00) |
| Module 21 (2274) | -0.16 (0.40) | 290 (0.25) | 8 (1.00) | 15 (1.00) |
| Module 22 (189) | -0.19 (0.30) | 26 (0.30) | 2 (0.47) | 4 (.71) |

**Note:** Column 1: module name and size (number of genes). Column 2: the correlation coefficient and associated *p*-value for correlation with behavioral state. Columns 3-4: the number of genes in the module that overlapped with the given gene list, and the associated *p*-value for these overlapping genes in Fisher's exact tests.

Two co-expression modules (Modules 4 and 7) were significantly correlated with behavioral state. Module 4 was enriched for genes in the Notch signaling pathway (Table S26), which has been demonstrated previously to mediate worker reproductive constraint in the presence of the queen (Duncan et al. 2016). Module 7 was not enriched for any GO terms or KEGG pathways, but it was overrepresented for pheromone binding proteins and signaling peptides with known roles in honey bee behavioral maturation, including vitellogenin, juvenile hormone esterase, and hexamerin 70a.

Six co-expression modules were significantly overrepresented for genes with parent-of-origin expression biases, including Modules 4 and 7 described above. Three of these additional modules (Modules 2, 3, and 8) were enriched for KEGG pathways and/or GO terms (Table S26). Module 2 was enriched for genes in the MAPK and Hippo signaling pathways, dorso-ventral axis formation, in addition to 109 GO terms for neural development, sensory perception, circadian rhythm, transcription regulation, and chromatin organization. Module 3 was enriched for genes in the Wnt signaling pathway, spliceosome, neuroactive ligand-receptor interactions, in addition to 76 GO terms for transcription regulation, ion transport, synaptic transmission, and neural development. Additionally, Module 8 was enriched for genes with protein binding function. Finally, three co-expression modules were enriched for genes predicted to be regulated by parent-biased transcription factors (Figure S6C), including Modules 2 and 3 described above, in addition to Module 10, which was enriched for genes related to nucleosome assembly and DNA-templated transcription (Table S26). Notably, Module 10 was also enriched for genes in our minimal support vector classifier of behavioral state (Table 1).

## Discussion

Here, we report evidence of a parent-of-origin effect on an altruistic behavior and associated physiological traits, and on transcription profiles of whole insect brains. Specifically, we found the proportion of worker bees within a colony responding to QMP was associated with the bees’ maternal lineage, as was the frequency of retinue response between individual bees. Ovary size and activation were also associated with maternal lineage, with bees that had larger and more active ovaries and were less responsive to QMP. In each reciprocal cross colony, bees had larger and more active ovaries when they came from crosses that were less responsive to QMP. Moreover, we found that maternally-biased transcripts were enriched in QMP-responsive worker bees relative to QMP-unresponsive bees. These results support a model in which bees with smaller ovaries—a trait which is determined during pre-adult development (Hartfelder et al. 2018)—are less likely to successfully compete with their sisters for ovary activation if the queen is lost (Makert et al. 2006) and are more likely to engage in the retinue response, increasing both their own and their sisters’ exposure to queen pheromone (Seeley 1979; Page 2013).

Together with previous studies, these results indicate that matrigenes favor the development of smaller ovaries while patrigenes favor the development of larger ovaries during pre-adult development. Likewise, matrigenes favor the retinue response while patrigenes inhibit it, and matrigenes inhibit ovary activation while patrigenes favor it (Kocher and Grozinger 2011; Galbraith et al. 2015; Kocher et al. 2015; Galbraith et al. 2016; Reid et al. 2017; Smith et al. 2020). Thus, during an individual worker’s life, patrigenes and matrigenes are in conflict over developmental, physiological, and behavioral traits.

Previous studies of the genetic basis of behavioral variation have revealed numerous insights into the coordination between genes, particularly within gene regulatory networks (GRNs). Behavioral stimuli are often associated with predictable and reproducible changes in brain gene expression profiles (Whitfield et al. 2003; Alter et al. 2008; Wong et al. 2015; Benowitz et al. 2019; Friedman et al. 2020; Kabelik et al. 2021; Werkhoven et al. 2021), which are controlled by GRNs (MacNeil and Walhout 2011). Thus, variation in the expression and regulation of genes within brain GRNs can be predictive of behavioral states (Sinha et al. 2020). In colonies of the honey bee (*Apis mellifera*), female worker bees exhibit extraordinary examples of behavioral variation between nestmates (Smith et al. 2022) and are tractable models for genetic experiments (Toth and Zayed 2021). Thus, studies in the honey bee have shown how transcriptional variation of genes within brain GRNs can contribute to variation in behavior (Chandrasekaran et al. 2011; Ingram et al. 2011; Mikheyev and Linksvayer 2015; Shpigler et al. 2017; Shpigler et al. 2019; Jones et al. 2020). Here, we explored how conflicts *within* these genes—which are predicted to occur due to parent-specific differences in optimal strategies for maximizing reproductive success (Gardner and Úbeda 2017)—can also contribute to behavioral variation.

Genes engaged in conflicts of origin “disagree” on how to use maternal resources (Gardner and Úbeda 2017). In mammals, this disagreement occurs within genes that regulate placental development and nutrient uptake by the fetus (Barlow and Bartolomei 2014). In plants, this conflict has been observed in genes that regulate nutrient deposition and storage in the endosperm of seeds, flowering time, fruit development, and root growth (Rodrigues and Zilberman 2015). In bumblebees, parent-biased genes in ovaries are associated with female gamete generation, regulation of ovulation, response to pheromone, and histone H3 acetylation (Marshall et al. 2020). Similarly, in honey bees, intragenomic conflict in worker ovary activation occurs in genes associated with neuron differentiation and embryonic development in fat body and ovaries (Galbraith et al. 2016), and in genes associated with reproduction-related hormonal signaling in brains (Galbraith et al. 2021). In our study, parent-biased transcripts in QMP-responsive bees were enriched for chromatin binding function, including several polycomb group genes (Pc, Sfmbt, RING1) and trithorax group genes (trx, Mi-2, polybromo, MTA3, Wdr82, Sbf), suggesting a potential role for chromatin regulation in mediating intragenomic in the retinue behavior.

Recent studies in plants and mammals have revealed a non-canonical epigenetic mechanism for genomic imprinting, which occurs through allelic differences in chromatin accessibility mediated by histone modifications, rather than allele-specific transcriptional repression by DNA methylation (DNAme) (Batista and Köhler 2020; Andergassen et al. 2021; Hanna and Kelsey 2021). In insects, except for paternal genome elimination, genomic imprinting appears absent (Coolon et al. 2012; Pegoraro et al. 2017), and DNAme does not repress transcription (Duncan et al. 2022). Thus, in honey bees and bumblebees, allele-specific differences in DNAme are not associated with allele-specific differences in transcription (Marshall et al. 2020; Wu et al. 2020). However, a mechanism of allele-specific transcriptional regulation like non-canonical imprinting has been described in *D. melanogaster* (Floc’hlay et al. 2021). Given that polycomb repressive complexes (PRCs) are more widely conserved than DNAme among organisms (Schuettengruber et al. 2017), analogous mechanisms of non-canonical imprinting may be present in insects. In honey bees, H3K27me by PRC2 appears to play a role in worker ovary activation (Duncan et al. 2020). Interestingly, PRC1 members *RING1*, which mediates H2AK119 ubiquitination (Tamburri et al. 2020), and *Pc*, which binds H3K27me3 to drive chromatin compaction and gene silencing (Erokhin et al. 2018), were both maternally-biased in QMP-responsive bees, whereas the H3K27 demethylase *Utx* (Smith et al. 2008) was paternally-biased. Taken together, these observations suggest that intragenomic conflict mediated by polycomb and trithorax group protein complexes may be a productive area for future studies in honey bees (Inoue 2023).

Several studies have demonstrated that transcriptional variation of co-regulated genes, versus large differences in individual genes, may be responsible for behavioral variation (Sinha et al. 2020). Additionally, meaningful transcriptomic differences associated with behavioral states may be missed by standard differential expression tests (Taylor et al. 2021). In our study, we did not detect any significantly differentially expressed genes between QMP-responsive and unresponsive bees, whereas a previous study of this behavior identified hundreds of differentially expressed genes (Kocher et al. 2010). The behavioral states of the bees we selected for sequencing (from one reciprocal cross per blocks 1, 3, and 4) were determined based on a single qualitative trait at a single timepoint rather than frequency of response to QMP across multiple timepoints (as in blocks 5 and 6 in our study and in Kocher et al. 2010). Thus, the transcriptomic differences associated with response to QMP in our study were likely too subtle to detect with standard differential expression tests. Therefore, we inferred co-expressed gene networks, identifying two modules that were correlated with behavioral state. Additionally, we identified a subset of the transcriptome that was predictive of behavioral state in a support vector machine (SVM) trained on the transcriptomic data, and the genes in this SVM were enriched in a co-expression module.

Our results provide support for the model of intragenomic conflict altering phenotypes by cascading impacts on downstream gene regulatory networks, versus directly altering RNA levels of a single gene with major effects (Patten et al. 2016). As in previous studies in bees (Galbraith et al. 2016; Marshall et al. 2020; Galbraith et al. 2021; Bresnahan et al. 2023), there was little overlap between genes that showed parent-biased expression and genes whose overall expression differences were associated with behavioral variation. However, our SVM and the combined set of genes showing parent-biased expression both showed overrepresentation for genes encoding proteins with DNA binding function, including several transcription factors. Moreover, we found that genes correlated with behavioral state, genes which predict response to QMP, and genes engaged in conflict, are enriched in conserved gene networks and functional pathways, and are potentially regulated by a set of transcription factors engaged in intragenomic conflict.

One of the co-expression modules that was correlated with behavioral state (Module 7) contained genes with known central functions in the retinue behavior and behavioral maturation, including vitellogenin (*Vg*) and juvenile hormone esterase (*JHe*). Oogenesis in insects involves the production of the egg yolk proteins from the egg yolk precursor Vg (Roy et al. 2018). In honey bees, Vg and juvenile hormone (JH) are coregulated in a double-repressor relationship and regulate behavioral maturation in workers: high Vg titers suppress JH and foraging behavior while high JH levels suppress Vg and nursing behavior (Harwood et al. 2017). In workers, QMP exposure suppresses JH biosynthesis, and JHe degrades JH (Mackert et al. 2008) delaying the transition from nursing to foraging (Pankiw et al. 1998).

The other co-expression module that was correlated with behavioral state (Module 4) was enriched for genes in the Notch signaling pathway. Chemical inhibition of Notch signaling in worker bees has been demonstrated to overcome the repressive effect of QMP on ovary activation (Duncan et al. 2016). Chemical inhibition of EZH2 (Tan et al. 2007), the enzymatic catalytic subunit of PRC2, has also been demonstrated to increase ovarian development in worker bees (Duncan et al. 2020). In our study, Module 4 was enriched for genes predicted to be regulated by the transcription factor BtbVII—which was maternally-biased in both QMP-responsive and unresponsive bees—including suppressor of hairless (*Su(H)*) and frizzled 2 (*fz2*). Su(H) carries the Notch receptor signal to the nucleus where it acts as a transcriptional activator of enhancer of split complex genes (Falo-Sanjuan and Bray 2020), suppressing neural development in *D. melanogaster* (Moore and Alexandre 2020). BtbVII was demonstrated to physically interact with Krüppel homolog 1 (Kr-h1) (Shokri et al. 2019), which is induced by JH and fz2 and regulates the expression of *Vg* (Jindra et al. 2015; Roy et al. 2018). Expression levels of *Kr-h1* and *fz2* in worker brains are downregulated by exposure to the queen or QMP (Grozinger and Robinson 2007) and are higher in foragers compared to nurse bees (Fussnecker and Grozinger 2008). Interestingly, Module 7 was also enriched for genes predicted to be regulated by a Notch signaling inhibitor, lethal (2) giant larvae (Langevin et al. 2005), which was paternally-biased in QMP-responsive workers.

One additional module was associated with response to QMP in that it was enriched for genes in our SVM classifier of behavioral state (Module 10). Both the SVM and module genes were overrepresented for structural constituents of chromatin and proteins with nucleosomal DNA binding function. This module was also enriched for genes predicted to be regulated by five parent-biased TFs, including fruitless (fru), scratch, stripe (referred to as early growth response protein 1 (Egr-1) in honey bees (Ugajin et al. 2013)), estrogen-related receptor (ERR), and runt. Egr-1 has been implicated in the nurse-to-forager transition in worker bees, potentially by rapidly inducing neural development (Lutz and Robinson 2013; Khamis et al. 2015; Shah et al. 2018). In *D. melanogaster*, both fru (Dalton et al. 2009) and ERR (Kovalenko et al. 2019) have been shown to regulate similar genes known to be induced by ecdysone receptor (EcR) signaling. In honey bees, *EcR* knockdown is associated with differences in transcript levels of several genes discussed above, including *Kr-h1* (Mello et al. 2014), and QMP exposure is associated with high ecdysteroid titers (Trawinski and Fahrbach 2018). In our study, *fru* was paternally-biased in unresponsive workers, whereas *ERR* was maternally-biased in responsive workers.

Taken together, the results of our study support predictions of the Kinship Theory of Intragenomic Conflict regarding the influence of maternally inherited genes on altruistic behaviors that increase an individual’s inclusive fitness. We report parent-of-origin effects on the retinue response behavior, ovarian development, and transcription in worker bee brains. We present evidence for intragenomic conflicts within conserved signaling pathways associated with behavioral maturation, neural development, and the regulation of reproduction in insects. Additionally, we suggest roles for transcription factors and chromatin regulation in parental allele-specific gene expression. Our study provides new insights into the genetic basis of social behavior and the potential for intragenomic conflicts as a driver of behavioral variation in honey bees and potentially other species.

## Data and Resource Availability

Scripts to reproduce the results of this study and detailed markdowns describing their use are available at https://github.com/sbresnahan/IGC-retinue. Sequencing reads for this study are deposited in the NCBI Sequence Read Archive under project accession PRJNA732718.

## Acknowledgements

We would like to thank Kaleena Cazajkowski (Penn State), E. T. Ash (Texas A&M) and Elizabeth (Liz) Walsh (Texas A&M) for assistance with creating and managing the honey bee stocks used in this study. We are grateful to members of Michael Axtell’s lab (Penn State) for feedback on our statistical methods, and members of the Grozinger lab for critical reading of the manuscript. This work was supported by the National Science Foundation (grant mo. MCB-0950896 to C.M.G.), as well as funding from the Huck Institute for the Life Sciences at Penn State. STB was supported by the National Science Foundation Graduate Research Fellowship Program (Grant # DGE1255832).

