## Supplementary material for "Beyond conflict: kinship theory of intragenomic conflict predicts individual variation in altruistic behavior": Dataset S4

**Dataset S4: figures for A Test of the Kinship Theory of Intragenomic Conflict in the Altruistic, Pheromone-Mediated Retinue Behavior in Honey Bees (Apis mellifera)**

Sean Bresnahan, Huck Institutes of the Life Sciences, Pennsylvania State University

2023-05-17

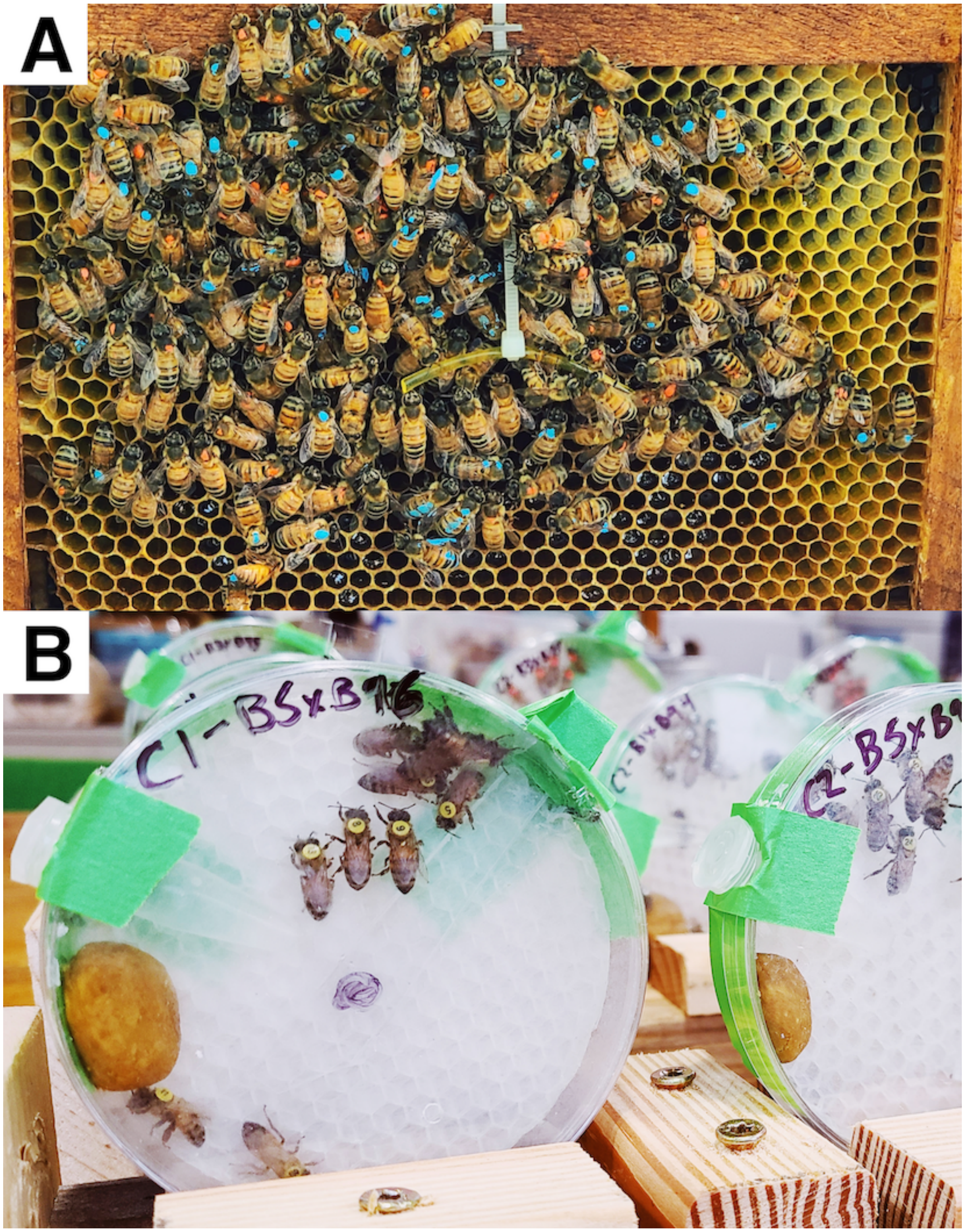


Figure S1: Retinue response assay. (A) Colony-level assay. Bees were marked with paint to identify their maternal lineage of origin and provided a strip of queen mandibular pheromone (QMP). (B) Individual-level assay. Bees were individually number tagged and provided fresh QMP before each assay via a removable slide.

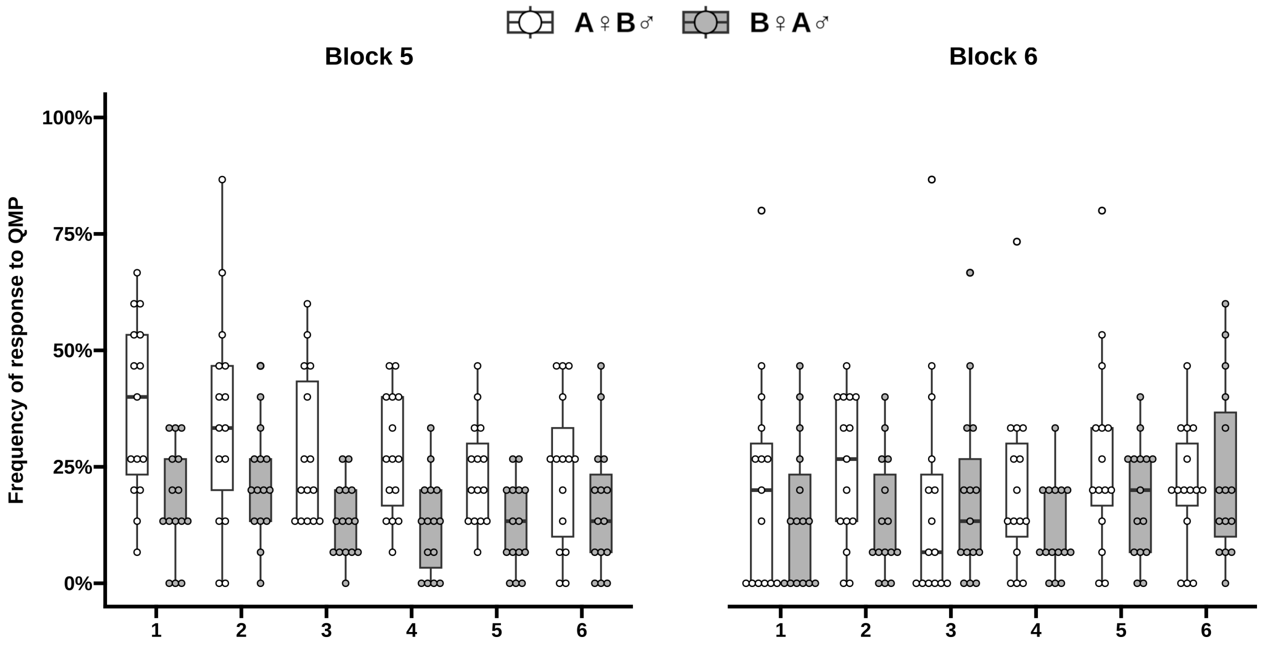


Figure S2: Retinue response of individual bees from each cross, separated by cage.

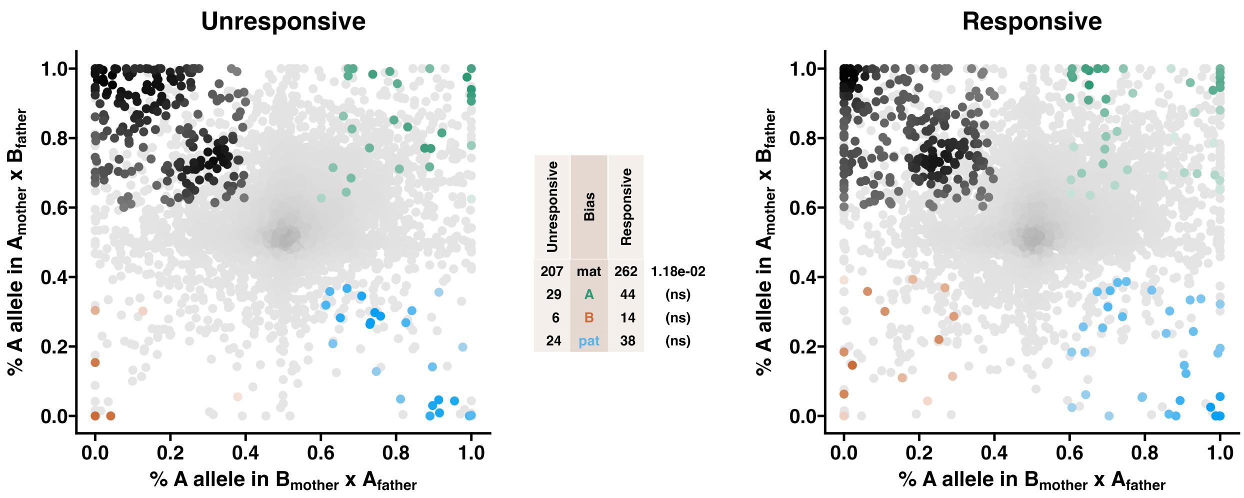


Figure S3: Response to QMP is associated with maternal allele-biased transcription. Allele-specific transcriptomes were assessed in worker bees from a reciprocal cross between different stocks of EHBs (C1xC3, Block 1) that were unresponsive or responsive to QMP. The x-axis represents, for each transcript, the proportion of cross A (C3 lineage) reads in bees with a cross B (C1 lineage) mother and cross A father (p1), and the y-axis represents, for each transcript, the proportion of cross A reads in bees with a cross A mother and cross B father (p2). Each color represents a transcript which is significantly biased at all tested SNP positions: black is maternal, purple is cross A, gold is cross B, blue is paternal, and gray is not significant. Significance was determined using the overlap between two statistical tests: a generalized linear interactive mixed model (GLIMMIX), and a Storer-Kim test along with previously established cutoff thresholds of $p1 < 0.4$ and $p2 > 0.6$ for maternal bias, $p1 > 0.6$ and $p2 < 0.4$ for paternal bias, $p1 < 0.4$ and $p2 < 0.4$ for lineage B bias, and $p1 > 0.6$ and $p2 > 0.6$ for lineage A bias.

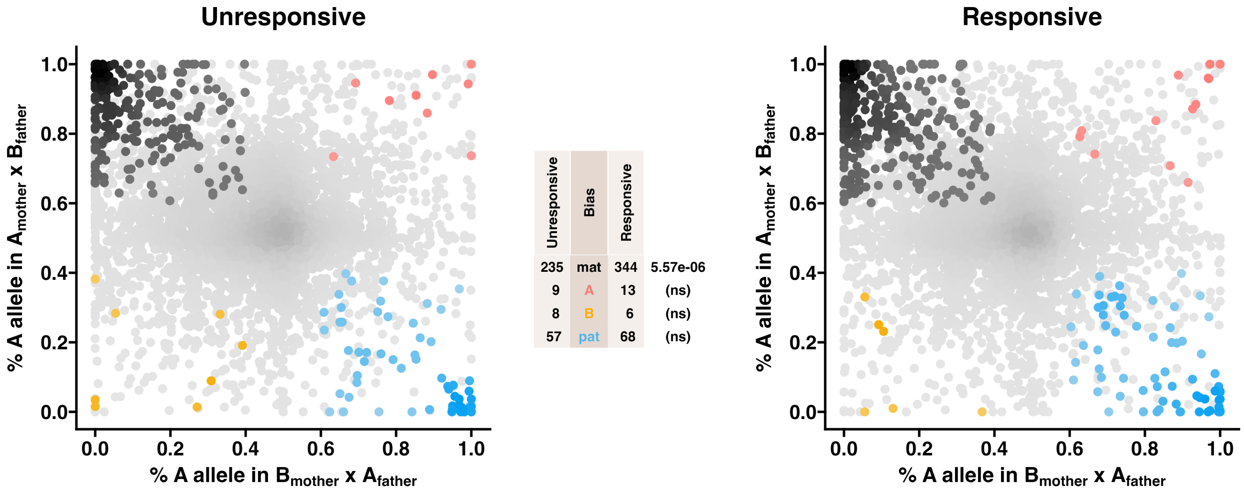


Figure S4: Response to QMP is associated with maternal allele-biased transcription. Allele-specific transcriptomes were assessed in worker bees from a reciprocal cross between different stocks of EHBs (C6xC11, Block 4) that were unresponsive or responsive to QMP. The x-axis represents, for each transcript, the proportion of cross A (C6 lineage) reads in bees with a cross B (C11 lineage) mother and cross A father (p1), and the y-axis represents, for each transcript, the proportion of cross A reads in bees with a cross A mother and cross B father (p2). Each color represents a transcript which is significantly biased at all tested SNP positions: black is maternal, purple is cross A, gold is cross B, blue is paternal, and gray is not significant. Significance was determined using the overlap between two statistical tests: a generalized linear interactive mixed model (GLIMMIX), and a Storer-Kim test along with previously established cutoff thresholds of $p1 < 0.4$ and $p2 > 0.6$ for maternal bias, $p1 > 0.6$ and $p2 < 0.4$ for paternal bias, $p1 < 0.4$ and $p2 < 0.4$ for lineage B bias, and $p1 > 0.6$ and $p2 > 0.6$ for lineage A bias.

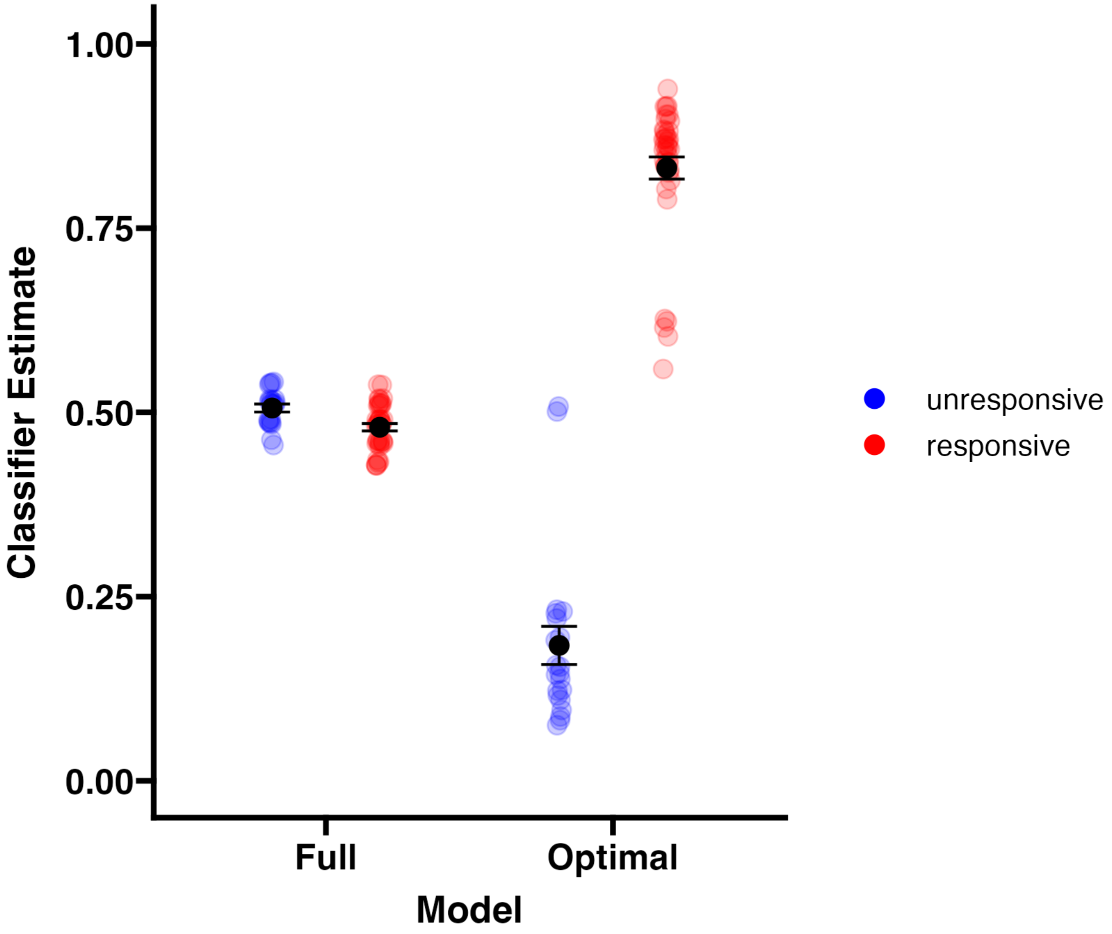


Figure S5: Support vector classifier estimates of test samples. Mean and standard error are indicated.

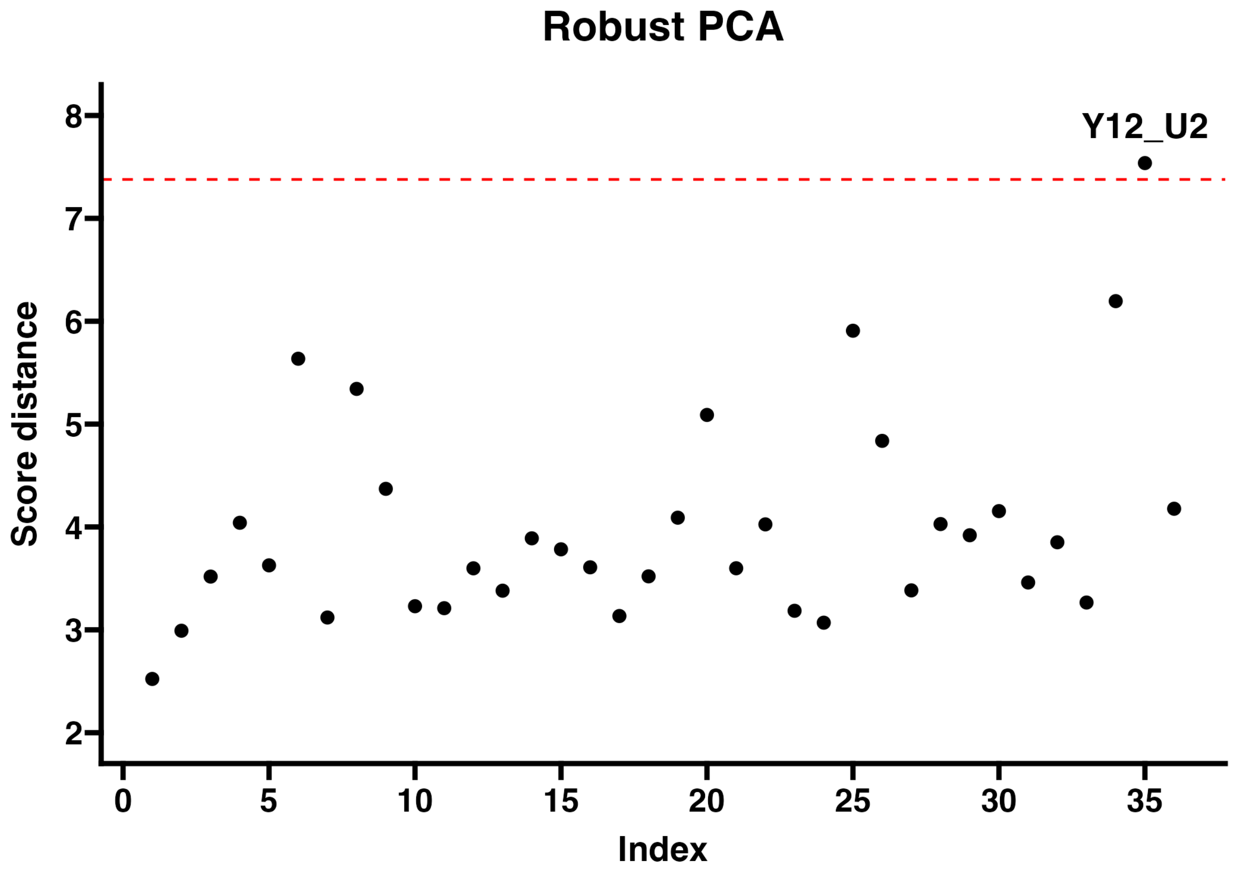


Figure S6: PCA projection pursuit method for outlier detection.

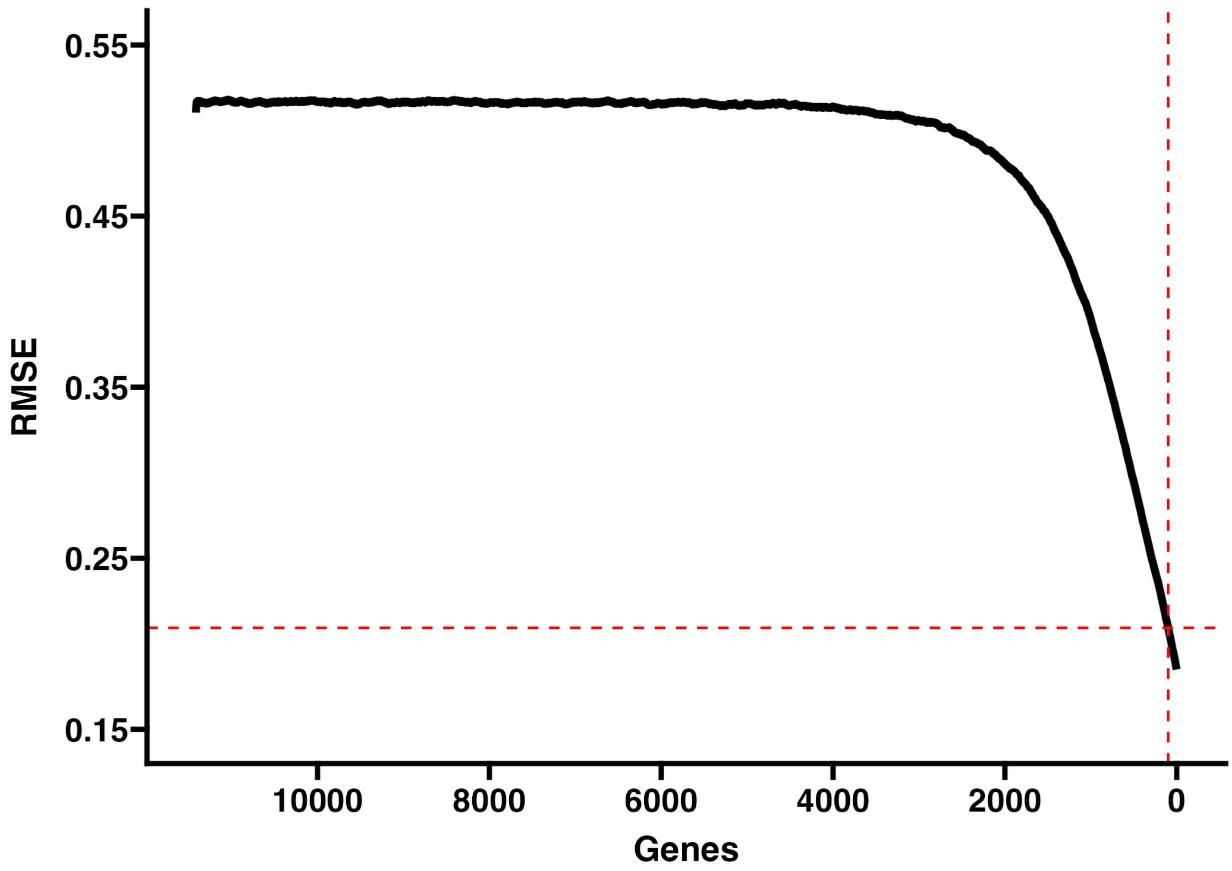


Figure S7: SVM classification error in feature selection.

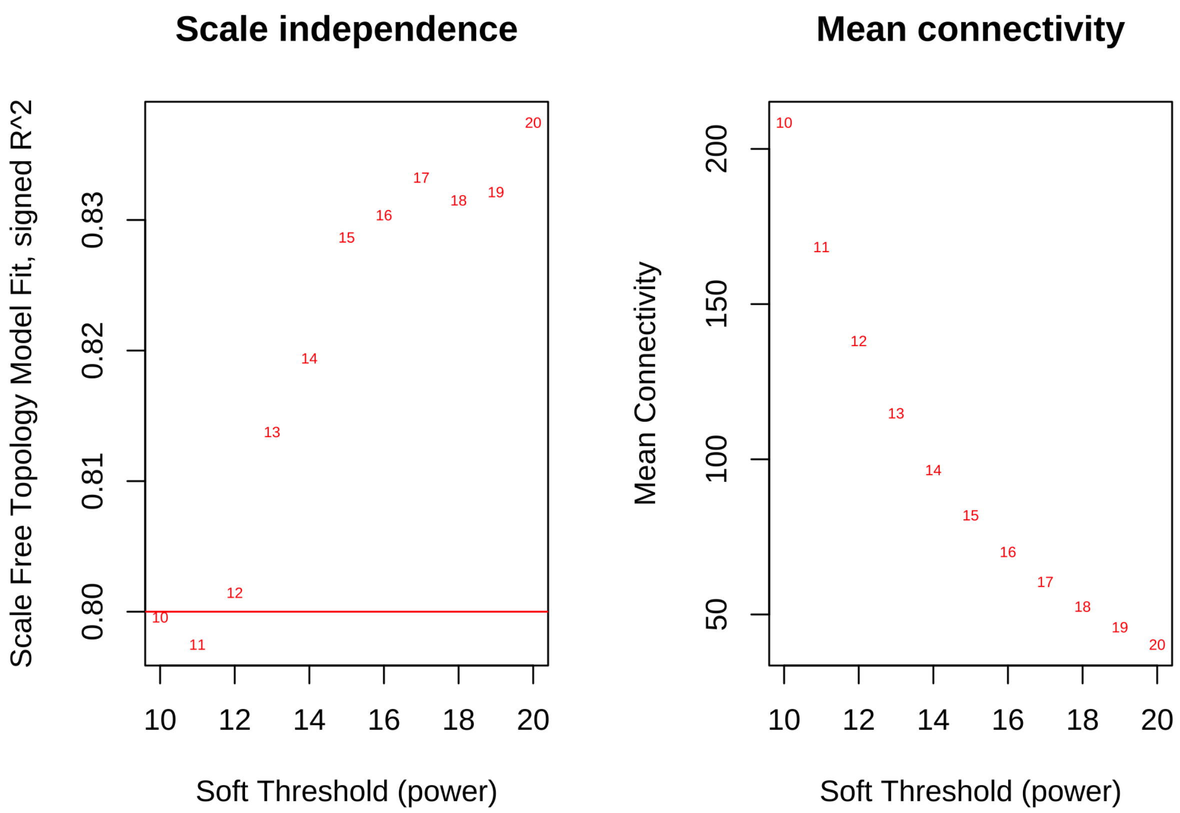


Figure S8: WGCNA diagnostic plots: scale independance and mean connectivity.

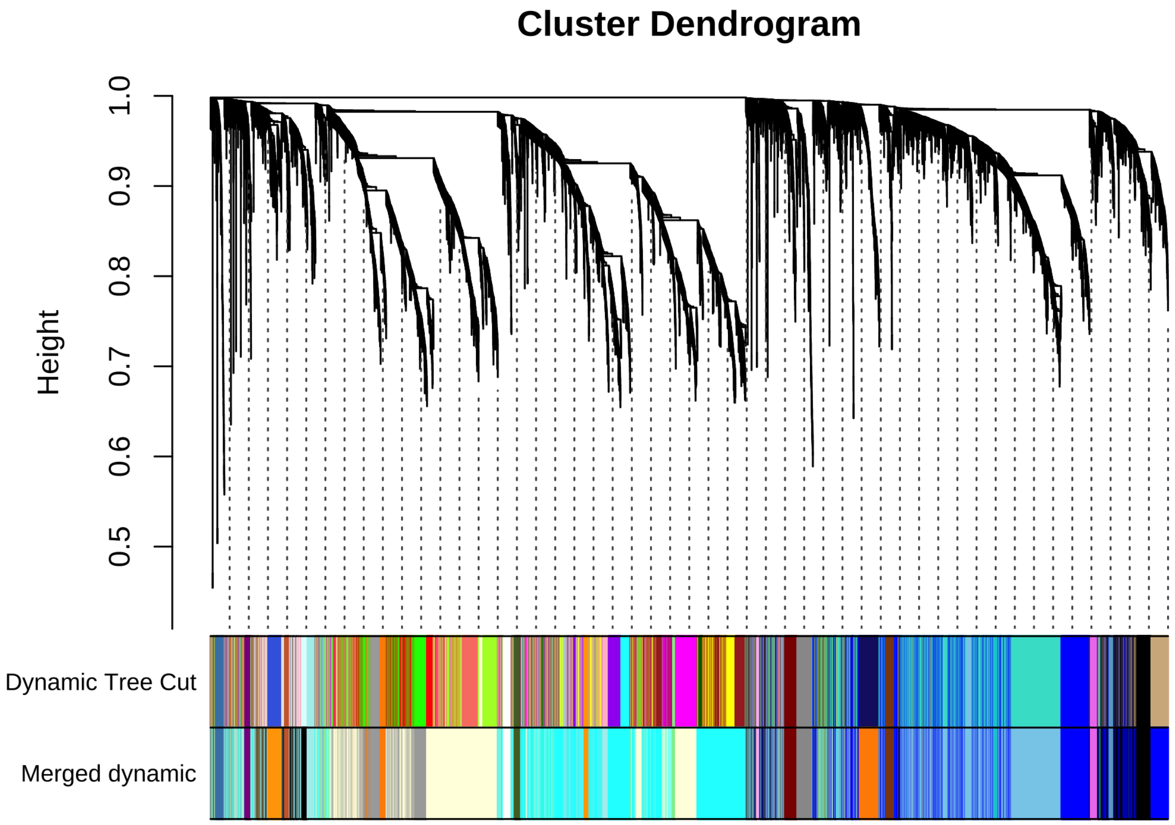


Figure S9: Gene coexpression network dendrogram. Modules with eigengene correlation coefficients > 0.75 were merged.
